# Explaining and predicting pseudoprogression in immunotherapy through joint modeling of immune infiltration and ctDNA dynamics

**DOI:** 10.64898/2026.09.18.752698

**Authors:** Aaron Li, Emil Lou, Kevin Leder, Jasmine Foo

**Affiliations:** School of Mathematics, University of Minnesota, Twin Cities, MN, USA; Masonic Cancer Center, University of Minnesota, Twin Cities, MN, USA; Division of Hematology, Oncology, and Transplantation, Department of Medicine, University of Minnesota, MN, USA; Department of Industrial and Systems Engineering, University of Minnesota, Twin Cities, MN, USA

## Abstract

Pseudoprogression in response to immune checkpoint inhibitor (ICI) therapy can be a complex challenge during treatment response evaluation. Erroneous classification of progression may result in stopping effective treatment for patients, while delayed confirmation of progression can prolong an ineffective treatment course. We develop a mechanistic mathematical model of tumor-immune dynamics under ICI treatment in which observed tumor volume includes contributions from both tumor burden and immune infiltration. We show analytically how pseudoprogression can arise from increased immune infiltration and proliferation as well as continued tumor growth during the “ramp-up” phase. The model also incorporates circulating tumor DNA (ctDNA) dynamics, which we demonstrate are able to closely fit existing longitudinal tumor and ctDNA data and extract realistic parameter ranges to generate representative *in silico* data. Using model-generated cohorts, we train a random forest (RF) classifier using longitudinal tumor and ctDNA measurements to distinguish pseudoprogression and true progression, and demonstrate that it improves response classification relative to the Response Evaluation Criteria in Solid Tumors (RECIST 1.1) and provides classification earlier than the immunotherapy-specific iRECIST, even in the presence of tumor measurement noise. This work predicts that paired longitudinal tumor-ctDNA measurements may enable earlier discrimination of true progression from pseudoprogression, and motivates collection of imaging and ctDNA data synchronously during ICI therapy, especially in the setting of suspected disease progression.

## 1 Introduction

Pseudoprogression (PsP) is a phenomenon in which tumor lesions initially appear enlarged on radiographic imaging after the start of treatment, but eventually regress and exhibit sustained treatment response. This phenomenon is most commonly observed during treatment with immunotherapies, such as immune checkpoint inhibitors (ICIs), which reactivate antitumor immune response by blocking pathways such as PD-1/PD-L1 [1, 2]. Clinical use of ICIs has rapidly expanded over the past decade, and they are standard of care for an increasing set of cancer types. Though the precise mechanisms of PsP are unknown, it is thought to arise due to the influx of immune cells into tumors triggered by these therapies, localized inflammation and edema, and delayed response to treatment, all of which could plausibly produce temporary increases in imaged tumor size [3]. A meta-analysis of 17 ICI studies observed a 6% pooled incidence rate of pseudoprogression across various solid tumor types [4]. However, in glioblastoma patients following resection and radiotherapy, pseudoprogression has also been observed with incidence rate estimates as high as 36% [5]. PsP poses a major challenge in clinical decision-making, since both withdrawing therapies from patients who would benefit and prolonging ineffective therapies can have serious negative consequences.

Clinical response evaluation guidelines such as the Response Evaluation Criteria in Solid Tumors (RECIST) 1.1 have traditionally been effective for cytotoxic chemotherapies and even targeted agents for which two-dimensional assessment of growth vs involution of tumors provided an accurate depiction of treatment response. However, in the case of immunotherapy and related pseudoprogression, this same approach may erroneously classify these patients as experiencing progressive disease, leading to premature cessation of treatment. An immunotherapy-specific guideline, iRECIST, was thus devised to address this issue by classifying the first observation of progressive disease as unconfirmed progressive disease (iUPD). In such cases, the guideline recommends continuing treatment until the next scheduled assessment (4–8 weeks later), ultimately classifying the outcome as confirmed progressive disease (iCPD) if tumor regression or resolution of related inflammation is not observed [6]. In a meta-analysis comparing RECIST 1.1 and iRECIST, Park et al. [7] found iRECIST would reclassify 3.9% of RECIST 1.1 progressive disease patients as having stable disease or partial/complete response, but had no impact on the overall response rate and disease control rate. For patients experiencing true progression of tumors, the use of iRECIST criteria could result in delays in pivoting to salvage therapy and added exposure to drug and financial toxicity.

While tumor volume assessments alone appear to be insufficient, the incorporation of biomarkers such as circulating tumor DNA (ctDNA) or Immunoscore-IC [8, 9] have demonstrated potential to help distinguish these response patterns. Lee et al. [10] found that in a cohort of 125 melanoma patients receiving anti-PD-1 treatment, 9/9 pseudoprogressors exhibited a favorable ctDNA profile (greater than 10-fold decrease from baseline to week 12) while only 2/20 progressors exhibited a favorable ctDNA profile. Guibert et al. [11] conducted a case study on two lung adenocarcinoma patients who exhibited pseudoprogression in response to ICI treatment. At day 60, the patients were classified as experiencing progressive disease by RECIST 1.1, but eventually experienced partial response upon follow-up at day 120. This response was preceded significantly by a dramatic decrease in ctDNA at day 30 in both patients. Bratman et al. [12] did not examine pseudoprogression in detail, but found that combining RECIST and ctDNA evaluation at cycle 3 (6-7 weeks) helped stratify risk groupings in solid tumor patients treated with pembrolizumab. A deep learning model was able to detect pseudoprogression from magnetic resonance imaging (MRI) features with 89% sensitivity, but only 62% specificity [13]. These observations suggest that ctDNA dynamics may provide further insights for discriminating pseudoprogression, but the scarcity of pseudoprogression data makes robust statistical analysis challenging. Testing this hypothesis would require longitudinal datasets in which both ctDNA and tumor radiographic assessments are collected simultaneously in patients until confirmation of response. To address this gap, here we develop a mechanistic mathematical model of tumor-immune response to ICIs to understand the sources of pseudoprogression and explore the use of ctDNA dynamics for distinguishing from true progression early during treatment.

Mechanistic mathematical modeling of pseudoprogression in ICI treatment is limited. Butner et al. [14] developed a partial differential equation model of tumor response to ICI treatment that was capable of exhibiting pseudoprogression due to one possible mechanism of delayed treatment effect. Scibilia et al. [15] compared four models of tumor response to ICIs and fit them to longitudinal patient tumor size data. One of these models was able to exhibit pseudoprogression dynamics incorporating another mechanism: delayed proliferation of immune cells. These prior works did not explicitly considered the contribution of immune infiltration to apparent tumor growth, a major hypothesized mechanism of PsP, nor did they explore the use of non-tumor biomarkers such as ctDNA for identifying this phenomenon.

A separate line of modeling efforts has emerged to describe ctDNA shedding from evolving tumor populations. Avanzini et al. [16] developed a mechanistic model of ctDNA shedding, focusing on detection of new or recurrent tumors, rather than early prediction of treatment response. Shrestha et al. [17] developed a coupled SDE-ODE model of tumor-immune dynamics and biomarker release, but do not explore dynamics under treatment. Khan et al. [18] used longitudinal liquid biopsy of carcinoembryonic antigen (CEA) and mathematical modeling to forecast time to treatment failure in colorectal cancer. However, under their model the CEA biomarker was assumed to be directly proportional to the tumor burden. As such, the model framework is unable to capture dynamics such as ctDNA peaks [19] or pseudoprogression. Ribba et al. [20] conducted an empirical exploration of tumor-ctDNA dynamics (but did not examine pseudoprogression) and provide a biexponential modeling framework. The authors describe their tumor-ctDNA modeling work as introductory and emphasize the need for improved mechanistic modeling. In prior work, we developed stochastic branching process models of tumor-ctDNA dynamics under targeted therapy, chemotherapy, and radiotherapy and used them to identify treatment-specific ctDNA biomarkers of partial or complete response, but did not examine immunotherapy or pseudoprogression [21, 22].

Here, we introduce a mechanistic model of tumor-immune dynamics under ICI treatment based on grounded biological principles. Our model includes both tumor and immune cells in the calculation of measured tumor size, and models ctDNA shedding under ICI treatment. We demonstrate that this model can be fitted to clinical tumor size and ctDNA trajectories. Using this model, we provide an analytic interpretation of progression, pseudoprogression, and response, and demonstrate how pseudoprogression naturally arises from simple, well-established biological mechanisms. We then compare analytic and clinical classifications of pseudoprogression, and investigate how clinical sampling times and thresholds influence the frequency with which tumor growth trajectories are clinically classified as pseudoprogression. Finally, using model-generated cohorts, we evaluate whether a random forest classifier, combining early tumor burden and ctDNA measurements, can improve discrimination between distinguish progression, pseudoprogression, and response/stable disease, relative to solely-imaging based response criteria (RECIST 1.1 and iRECIST). This work suggests that prospective, synchronized longitudinal imaging and ctDNA measurements could improve response classification, and warrant further consideration as a strategy for distinguishing pseudoprogression from true progression.

## 2 Results

### 2.1 Model

We present an ordinary differential equation (ODE) model of tumor-immune and ctDNA dynamics under ICI treatment. Our model (Fig. 1) consists of tumor cells (*T*), active immune cells (*I*^*A*^), exhausted immune cells (*I*^*E*^), and ctDNA (*C*). The differential equations governing the system are given below.

**Figure 1.**
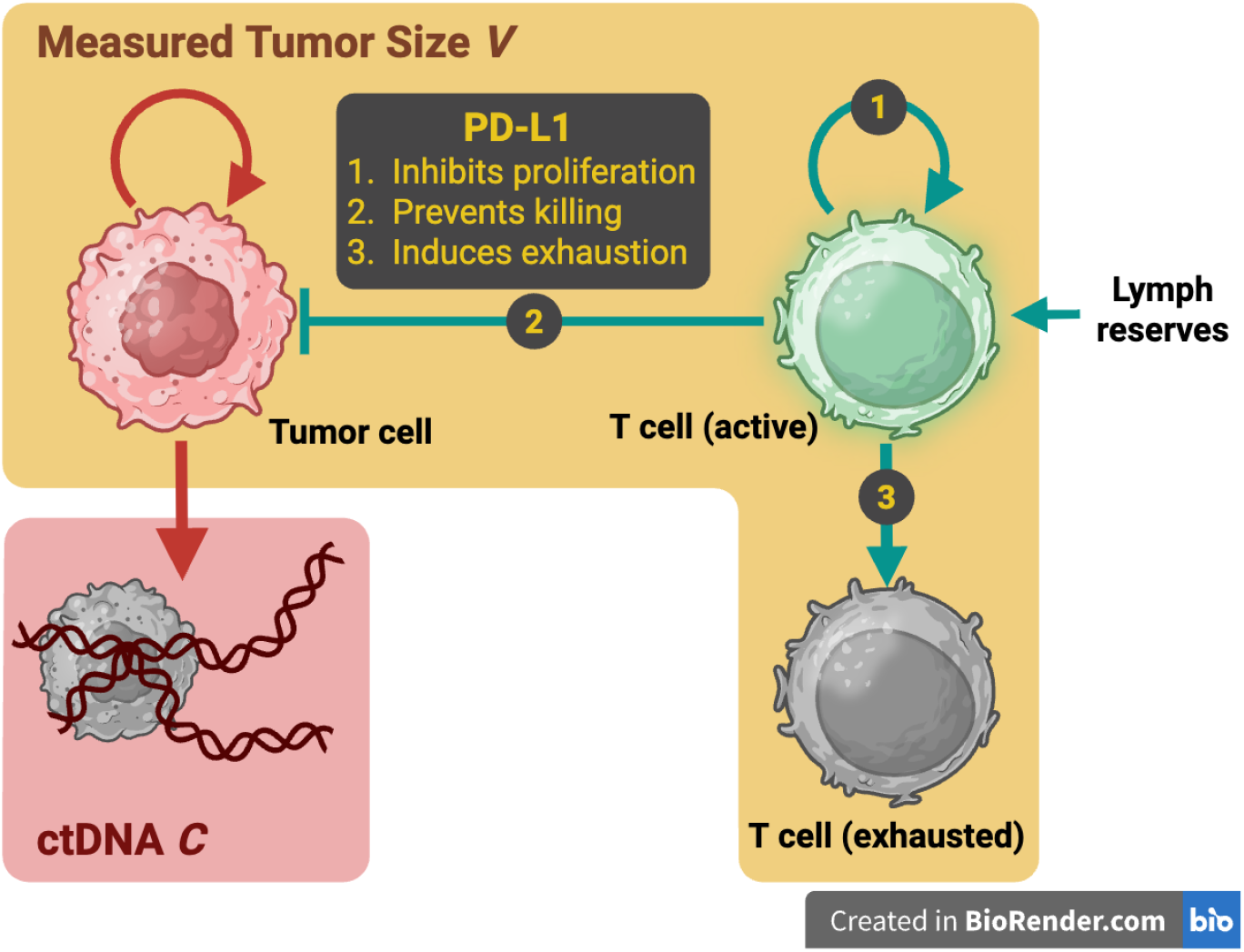
A schematic of our mechanistic model. Created in BioRender.

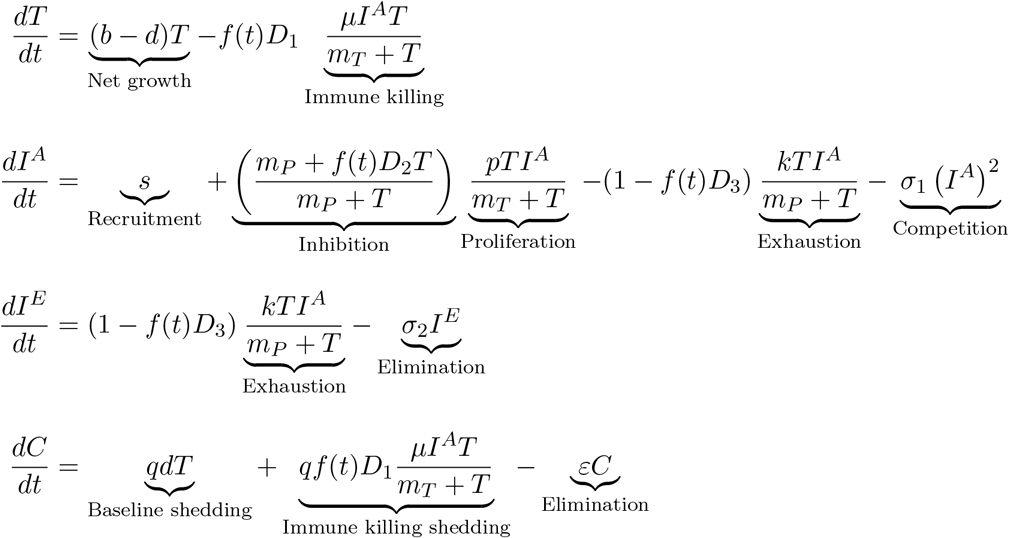

Many of the interactions in these equations are modeled by terms of the form

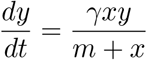

to represent *y* being stimulated or eliminated by interactions with *x*. The denominator provides a rate-limiting effect so that for large *x* the rate does not explode unrealistically. This Michaelis-Menten-like structure is well-established in tumor-immune modeling and is included in fundamental models such as that of Kirschner and Panetta [23].

#### Tumor dynamics

Since we are focusing on categorizing early response to treatment, exponential tumor growth is assumed, with base net growth rate *b* − *d* [24]. The immune killing term represents the rate at which tumor cells are eliminated by T cells. This term follows the rate-limited structure discussed above, reflecting a saturation effect in killing governed by parameter *m*_*T*_.

#### Immune dynamics

Baseline recruitment of active T cells from lymph node reservoirs is represented by *s*, as in Kuznetsov et al. [25]. The proliferation of the immune cells is stimulated by the presence of tumor cells, with *m*_*T*_ again representing rate-limitation due to spatial saturation. However, the presence of PD-L1 expression can inhibit T cell proliferation [26]. We model this by using the term 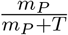 to downregulate the immune proliferation rate in the presence of PD-L1 expression. *m*_*p*_ represents the saturation of the antigen signal on the immune cells. Notice that when *T* = 0, this term is 1, representing no inhibition of T cell expansion. As *T* → ∞, this term goes to 0, representing full inhibition of T cell proliferation.

The PD-L1/PD-1 checkpoint can also lead to T-cell exhaustion [27]. The exhaustion term 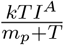 represents the transition from active immune cell *I*^*A*^ to exhausted immune cell *I*^*E*^, and follows a similar structure using *m*_*p*_ to model saturation of the PD-L1 signal. Finally, T cells can exhibit a rapid contraction [28] that may be influenced by self-competition or fratricide via Fas–FasL interaction [29, 30]. We include the *σ*_1_(*I*^*A*^)^2^ term to capture the density-dependent death rate.

#### ICI treatment

The drug concentration at time *t* will be given by *f* (*t*) ∈ {0, 1}. We omit more nuanced pharmacokinetic modeling for simplicity, and note that many ICI treatments, such as atezolizumab, exhibit flat response across a range of concentrations [31]. Without treatment (*f* (*t*) = 0), the tumor is assumed to completely escape immune-killing via the PD-L1/PD-1 checkpoint [2]. Tumor cells that express PD-L1 can inhibit immune killing, inhibit immune proliferation, and trigger immune exhaustion. We use *D*_1_, *D*_2_, and *D*_3_ to represent the ICI treatment’s efficacy at disrupting each of these effects, respectively. For example, examining the term representing inhibition of immune proliferation, we use *f* (*t*)*D*_2_ to represent the proportion of T cells that are shielded from the PD-L1 suppression, so

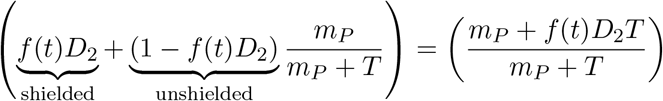

Notice that increasing *f* (*t*)*D*_2_ decreases the inhibitory effect, so the treatment can enable the proliferative “burst” of T cells observed *in vivo* [28]. Exhausted T cells are generally not able to be rescued by ICI treatment, and recovery is instead driven by peripheral T cells that have not been exhausted [2]. Thus, we do not include transitions from *I*^*E*^ to *I*^*A*^. Instead, exhausted immune cells simply decay at rate *σ*_2_.

The efficacy parameters must satisfy *D*_*i*_ ∈ [0, 1]. For *D*_1_, this ensures that the ICI improves immune killing but the effect cannot go beyond the immune system’s intrinsic maximum killing rate. We interpret *f*(*t*)*D*_2_, *f*(*t*)*D*_3_ as the fraction of cells that are shielded from checkpoint-mediated suppression of proliferation and from checkpoint-mediated exhaustion, respectively, so *D*_2_, *D*_3_ ∈ [0, 1].

#### Circulating tumor DNA dynamics

Similarly to a mechanistic model developed by Avanzini et al. [16], our model assumes that for each cell death, 1 human genome equivalent (hGE) of ctDNA is shed with probability *q* and the ctDNA breaks down and is eliminated from the bloodstream at rate *ε*. Note that the shedding includes both tumor cells that die from natural turnover as well as tumor cells that are eliminated by T cells.

#### Observed tumor volume

While existing models generally assume that measured tumor volumes correspond purely to tumor cell populations, the infiltration of immune cells into the tumor may increase its apparent volume [32]. Thus, we make the assumption that radiographically observed tumor volume reflects contributions from both tumor and intratumoral immune populations, and define the observed tumor volume as

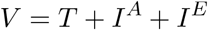

to incorporate the contribution of active and exhausted T cells to the measured tumor volume. We approximate the cells as being equal in size as they are within an order of magnitude [2]. Moreover, a parameter introduced to account for any relative difference in size could be absorbed by performing a parameter reduction to an equivalent model and simply lead to a minor scaling adjustment of parameter values.

Let *V*_0_ and *C*_0_ be the observed tumor size and ctDNA quantity at the start of treatment, respectively. Since longitudinal tumor size and ctDNA data are often presented in terms of relative changes [12], we can normalize our system by performing the following parameter reduction:

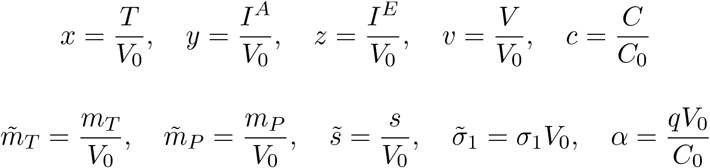

Moreover, focusing on the on-treatment dynamics, we can fix *f* (*t*) = 1 and further reduce the parameter space by setting 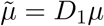 and 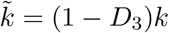. This results in the system below:

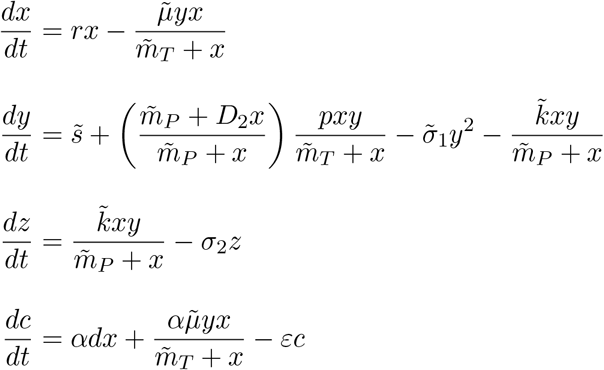

### 2.2 Model fitting to clinical data

We next investigated whether the model can reproduce clinically observed tumor dynamics. Similarly to Scibilia et al. [15], we evaluated our model using four sets of longitudinal tumor data from non-small cell lung cancer (NSCLC) patients treated with atezolizumab: POPLAR [33], BIRCH [34], OAK [35], and FIR [36]. After pooling the data and filtering for patients with at least four longitudinal tumor measurements, the dataset consisted of *n* = 632 patients. We used non-linear least squares via `scipy.optimize.least_squares` to fit *V* = *x* + *y* + *z* to the patient trajectories, using 10 random restarts per patient and an additional 70 restarts for patients with *R*^2^ *<* 0.80. The model closely reproduced observed trajectories, achieving a median *R*^2^ = 0.9982 and a median normalized mean absolute error (MAE) of 0.006, which is equivalent to a median MAE of 0.07 cm^3^. This demonstrates the model’s ability to reproduced observed trajectories, but does not uniquely identify underlying biological parameters. Indeed the number of parameters in our model relative to the low number of data points available for many patients. However, the purpose of this fitting is not to infer unique patient-specific biological parameters, but rather to establish that the model can reproduce clinically observed tumor dynamics and to inform parameter distributions for subsequent analyses. Fig. 2 shows example fits to tumor data from the BIRCH trial [34] for patients exhibiting progression, pseudoprogression, and disease response. In order to increase parsimony, we fixed *D*_2_ = 0.75, *σ*_2_ = 0.03, and *m*_*P*_ = *V*_0_*/*10^6^ at their cross-patient medians because they exhibited the lowest variance among parameters and fixing them did not result in decreased quality of fits. The values for the other parameters are summarized in Table 1.

**Table 1.** Parameter ranges resulting from fitting model to pooled longitudinal tumor dataset of *n* = 632 NSCLC patients treated with atezolizumab. Here 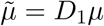 and 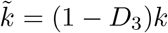 denote the net immune killing and net immune exhaustion rates on treatment. From these values we derive *y*_0_ = (1 − *x*_0_)*y*_frac_ and *z*_0_ = 1 − *x*_0_ − *y*_0_.

| Parameter | Mean ± Standard Deviation | Minimum | Maximum | Units |
| --- | --- | --- | --- | --- |
| $r$ | $0.018 \pm 0.027$ | $1.0 \times 10^{-4}$ | 0.10 | $\text{day}^{-1}$ |
| $\tilde{\mu}$ | $0.392 \pm 0.273$ | 0.005 | 1.00 | $\text{day}^{-1}$ |
| $\tilde{m}_T$ | $0.217 \pm 0.305$ | $1.0 \times 10^{-6}$ | 1.00 | — |
| $\tilde{s}$ | $0.0019 \pm 0.0032$ | $1.0 \times 10^{-8}$ | 0.010 | $\text{day}^{-1}$ |
| $p$ | $0.343 \pm 0.267$ | 0.010 | 1.00 | $\text{day}^{-1}$ |
| $\tilde{\sigma}_1$ | $3.12 \pm 7.65$ | $1.0 \times 10^{-3}$ | 50.0 | $\text{day}^{-1}$ |
| $\tilde{k}$ | $0.108 \pm 0.089$ | $2.5 \times 10^{-4}$ | 0.25 | $\text{day}^{-1}$ |
| $x_0$ | $0.758 \pm 0.179$ | 0.50 | 1.00 | — |
| $y_{\text{frac}}$ | $0.394 \pm 0.337$ | 0 | 1.00 | — |

**Figure 2.**
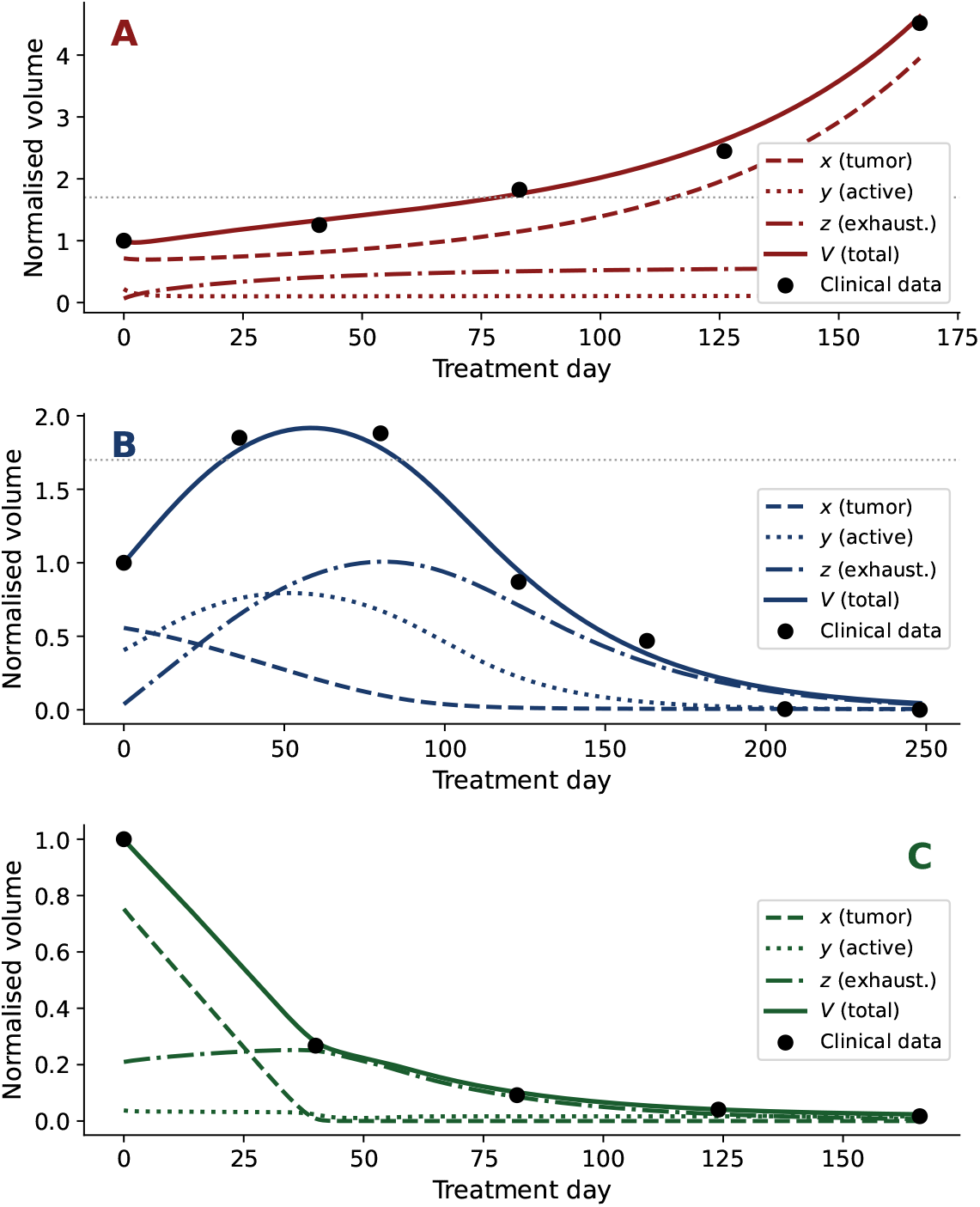
Example clinical trajectories of from the BIRCH trial [34] along with our ODE model fitting exhibiting (a) progression (*R*^2^ = 0.996), (b) pseudoprogression (*R*^2^ = 0.993), and (c) response (*R*^2^ = 0.999).

Bratman et al. [12] collected longitudinal tumor and ctDNA samples in solid tumor patients treated with pembrolizumab. *n* = 23 patients had at least 4 tumor measurements and at least 2 ctDNA measurements (including baseline). Fitting our model to this cohort produced a median *R*^2^ = 0.98 for tumor size fitting and a median *R*^2^ = 0.92 for the ctDNA fitting.

### 2.3 Defining pseudoprogression

#### 2.3.1 Analytic definition of pseudoprogression

We will now show how pseudoprogression dynamics can naturally arise from the biological mechanisms represented in the our model. Let us focus on tumor cells (*x*) and active immune cells (*y*), the active components of the system, and let the treatment be fixed at *f* (*t*) = 1. This scenario represents a simplified version of conditions at the start of treatment. We use a simplified model in this analytic context to explore the mechanistic behaviors driving pseudoprogression. Let *n*_*x*+*y*_ denote the nullcline of the system, the curve for which 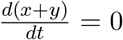 (Section 4.1 and Appendix A). Let *S* represent the separatrix of the system beyond which *x* → ∞. Finding a closed form of *S* for this nonlinear system is not tractable, but we can derive numerical solutions.

We now define the analytic response categories in terms of the position of the initial condition (*x*_0_, *y*_0_) relative to *n*_*x*+*y*_ and *S*.

- **Progression:** *y*_0_ < *S*(*x*_0_) Trajectories that begin below the separatrix will result in *x* → ∞, corresponding to tumor escape and disease progression. These trajectories will be described as **analytic progressors (aP)** (Fig. 3A).
- **Pseudoprogression:** *y*_0_ > *S*(*x*_0_) and *y*_0_ < *n*_*x*+*y*_(*x*_0_) Trajectories beginning in this region will result in eventual disease control or eradication because the initial condition (*x*_0_, *y*_0_) is above the separatrix. Since (*x*_0_, *y*_0_) is below the nullcline *n*_*x*+*y*_, *x* + *y* will be increasing initially. Thus, points in this region will result in initial transient increases, followed by response or stable disease. We will call these trajectories **analytic pseudoprogressors (aPsP)**. Fig. 3B shows an example aPsP trajectory. Notice that the apparent increase in *V* = *x* + *y* is driven by both mechanisms: delayed immune activity allowing continued tumor growth initially as well as increased volume as a result of immune infiltration.
- **Response/stable disease:** *y*_0_ > *S*(*x*_0_) and *y*_0_ > *n*_*x*+*y*_(*x*_0_) Trajectories that begin above the separatrix and the nullcline will immediately decrease and maintain response or stable disease in the long term and will be considered **analytic response/stable disease (aRSD)** (Fig. 3C).

**Figure 3.**
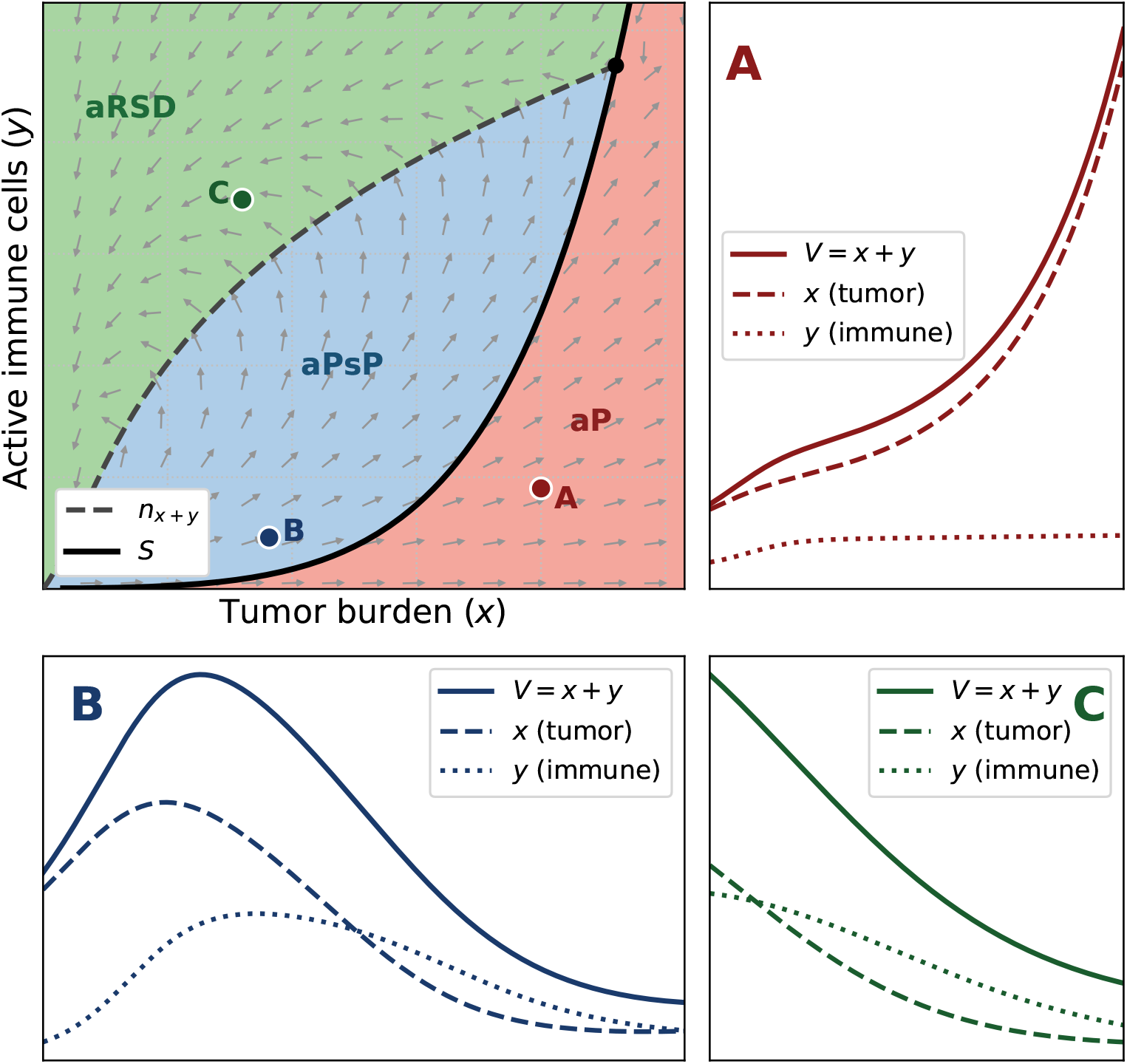
An example phase plane of the tumor burden *x* and the active immune cells *y*. Example trajectories for (a) analytic progression, (b) analytic pseudoprogression, and (c) analytic response/stable disease. The corresponding initial conditions are indicated on the phase plane.

Fig. 3 shows an example phase plane of *x* and *y* with the nullcline and separatrix plotted and the aP (red), aPsP (blue), and aRSD (green) shaded. Example trajectories are also shown with their initial conditions indicated on the phase plane. This analytic framework demonstrates how pseudoprogression arises naturally from the underlying biological mechanisms in our model. While the separatrix distinguishes potential trajectories based on their end behavior, the position of the initial conditions relative to the nullcline determines whether the total volume will increase or decrease initially. Continued tumor growth and immune infiltration can both contribute to an initial increase in volume.

#### 2.3.2 Comparison with clinical pseudoprogression

We now return to our full model, which includes exhausted immune cells. Using the parameter values obtained from fitting (Table 1), we generated an *in silico* cohort of 10,000 patients with longitudinal tumor data. For each patient, we simulated the collection of tumor measurements at baseline and every 25–50 days after the initiation of treatment, up to day 300. Each measurement for the cohort examined here included tumor diameter measurement noise with 10% coefficient of variation.

Following iRECIST [6], we define a clinical classification of each trajectory as follows:

- Disease progression (cP): A sustained increase in the sum of long diameters (SLD) of at least 20% relative to baseline (equivalent to a 70% increase in volume).
- Pseudoprogression (cPsP): At least 20% increase in SLD, followed by stable or responsive disease.
- Response/stable disease (cRSD): Trajectories that do not cross the 20% increase threshold at all.

Since this categorization requires a follow-up sample to distinguish cP and cPsP, 66 patients who first exhibited progression directly before the day 300 time horizon and did not receive a follow up measurement were excluded, resulting in an *in silico* cohort of *n* = 9,934 patients.

Fig. 4 shows a confusion matrix for the analytic and clinical classifications. Note that concordance between the two classifications is generally good, other than the large group of trajectories classified as PsP analytically, but not clinically. Further investigation reveals that this behavior is expected because many trajectories that exhibit a transient increase will not meet the clinical threshold of a 70% increase in volume (20% increase in the sum of long diameters) required to qualify as PsP. Of the 4,105 trajectories in this group, 7.1% did increase by at least 70% between sample times (PsP missed due to sampling), 75.7% exhibited a transient increase but did not exceed a 70% increase (PsP too small to qualify), and 17.2% were classified as responders because the elimination of exhausted immune cells (which do not factor into the analytic classification) obscured the PsP dynamics completely.

**Figure 4.**
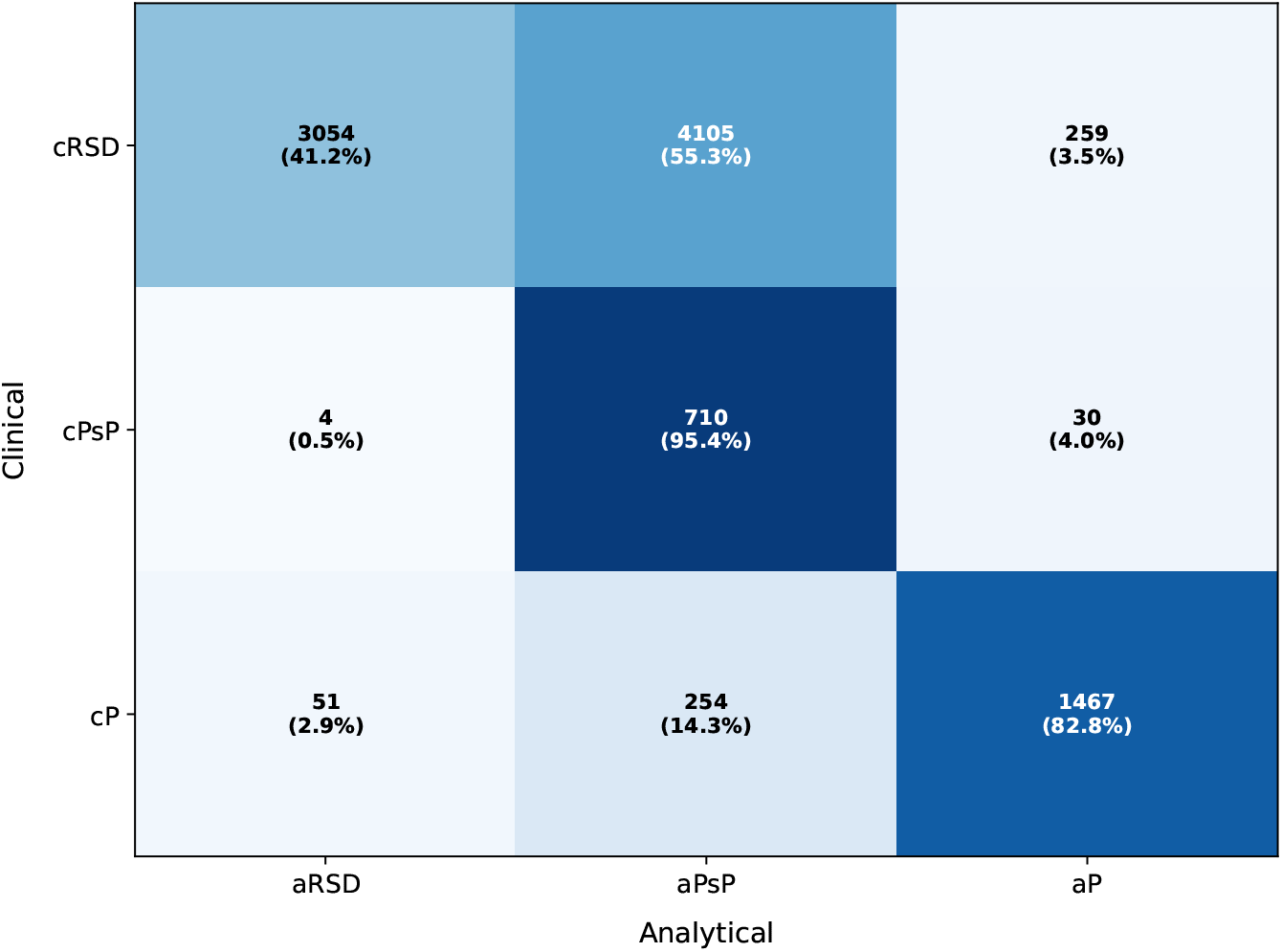
A confusion matrix for the analytic and clinical classifications of the *n* = 9,934 *in silico* patients as progression (P), pseudoprogression (PsP), and disease response/stable disease (RSD). The data for this cohort was sampled with a clinical schedule and included 10% tumor diameter measurement noise and 30% ctDNA measurement noise. The off-diagonal aPsP/cRSD group is expected given that the clinical classification of PsP relies on discrete time samples and requires growth to surpass a threshold of 70% increase in volume.

The effects of the clinical threshold and sampling interval are further examined in Fig. 5. The heatmap shows the simulated prevalence of pseudoprogression in the cohort as the threshold for detection of progression varies from 1.05 to 2.0 and the sample interval varies from 1 day to 75 days. Increasing the threshold requires a more pronounced transient peak for PsP classification while increasing the sampling interval increases the likelihood that a peak is missed between samples. Varying these conditions in this range resulted in PsP prevalence ranging from 4.2% at the most restrictive and 24.2% at the most broad.

**Figure 5.**
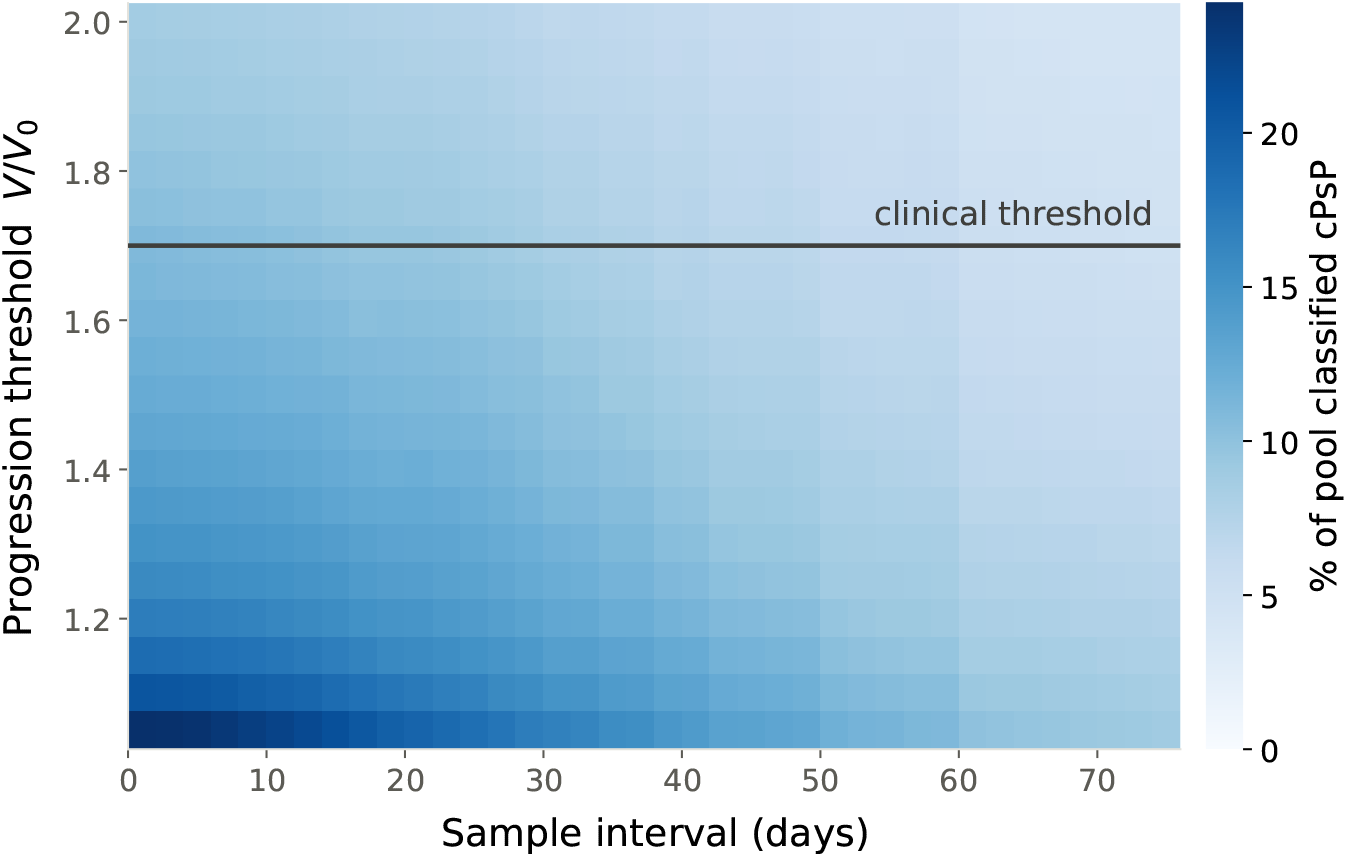
Clinically, pseudoprogression generally requires an observed increase of at least 70% in tumor volume (approximately 20% increase in the sum of long diameters), followed by tumor control or response. The volume increase threshold and the length of the interval between samples can both influence whether a trajectory that exhibits a transient increase will be categorized as pseudoprogression. The heatmap shows the percentage of the *n* = 9,934 patients that would be categorized as PsP for *V/V*_0_ thresholds between 1.05 and 2.0 and sample intervals from 1 day to 75 days. At *V/V*_0_ = 1.05 with a sample interval of 1 day, the rate of PsP was 24.2% and at a threshold of 2.0 with intervals of 75d, the rate of PsP was 4.2%.

### 2.4 Prediction and classification

In order to assess the potential predictive potential of ctDNA sampling, we augment our *in silico* cohort to include ctDNA trajectories as well. Using the parameter values described in Table 2, we simulated the collection of ctDNA samples every 25–50 days to accompany the tumor measurements.

**Table 2.** Parameters used for generation of ctDNA trajectories.

| Parameter | Definition | Units |
| --- | --- | --- |
| $\phi$ | $d/b$ ; $\phi \sim \text{Uniform}(0.20, 0.90)$ | — |
| $\varepsilon$ | 25 (fixed; plasma half-life $\approx 40$ min [37]) | $\text{day}^{-1}$ |
| $d$ | $r\phi / (1 - \phi)$ | $\text{day}^{-1}$ |
| $\alpha$ | $(\varepsilon + r) / (d x_0) = (1 - \phi)(\varepsilon + r) / (r\phi x_0)$ | — |

Using our *in silico* cohorts, we developed random forest (RF) classifiers and assessed their ability to accurately predict patient response. The models were evaluated on their ability to:

1. Distinguish between progression, pseudoprogression, and disease response (“3-class” categorization).
2. Detect whether observed progression is true progression or pseudoprogression.

To quantify performance on these tasks, we used Cohen’s Kappa [38], a statistical measure of inter-rater reliability for categorical data that is suitable for imbalanced data sets. The ground truth classification for each patient was determined using the full trajectory of longitudinal tumor volume samples every 25-50 days until day 300.

#### 2.4.1 Coupled tumor-ctDNA data improves classification performance

First, we compared three random forest classifiers: one with access to tumor and ctDNA data, one with ctDNA only, and one with tumor only. These three RF models were evaluated on the clean, no noise cohort as well as a cohort with 10% tumor diameter measurement noise and 30% ctDNA measurement noise. As shown in Fig. 6, the classifier with access to both tumor and ctDNA data, RF (all), outperformed the RFs based on ctDNA and tumor data alone in both tasks and both cohorts. In the subsequent sections, RF will refer to RF (all) unless otherwise specified.

**Figure 6.**
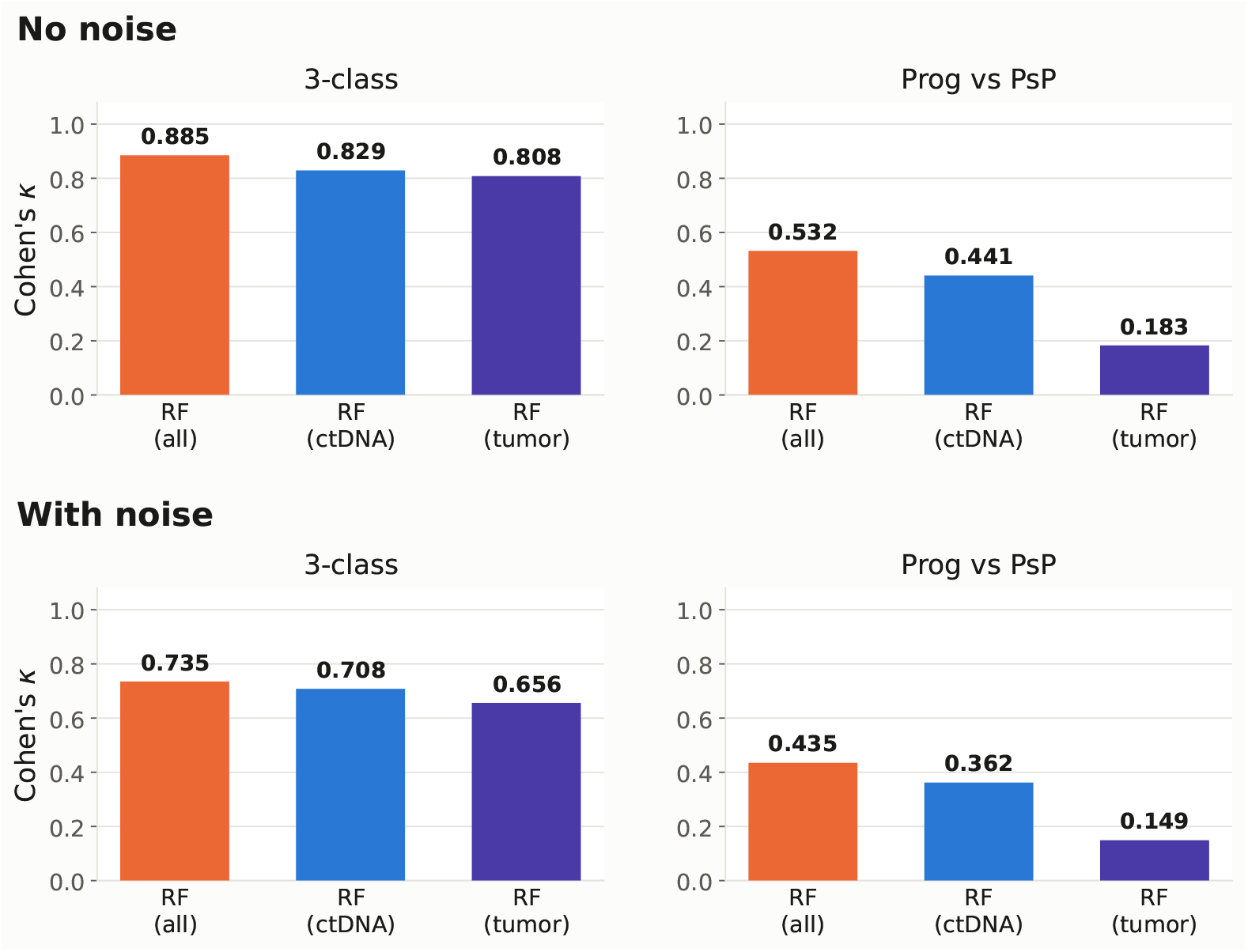
Cohen’s Kappa for classification of *n* = 9,934 *in silico* trajectories by random forest models with access to tumor and ctDNA data, ctDNA only, and tumor only. Performance is analyzed for the 3 category classification with progression, pseudoprogression, and response/stable disease as well as for distinguishing progression and pseudoprogression. The models are compared on data from the clean cohort as well as data from a cohort with 10% tumor SLD noise and 30% ctDNA noise applied.

#### 2.4.2 RF performance robust to ctDNA noise and sampling interval

Next, we evaluated the performance of the random forest classifier with varied levels of lognormal ctDNA measurement noise (Fig. 7). For this evaluation, 10% tumor diameter measurement noise was added for each scenario. As the coefficient of variation ranges from 0-40%, the *κ* for the RF classifier remains steady for the 3-class task, and diminishes only slightly in the progression vs. pseudoprogression task.

**Figure 7.**
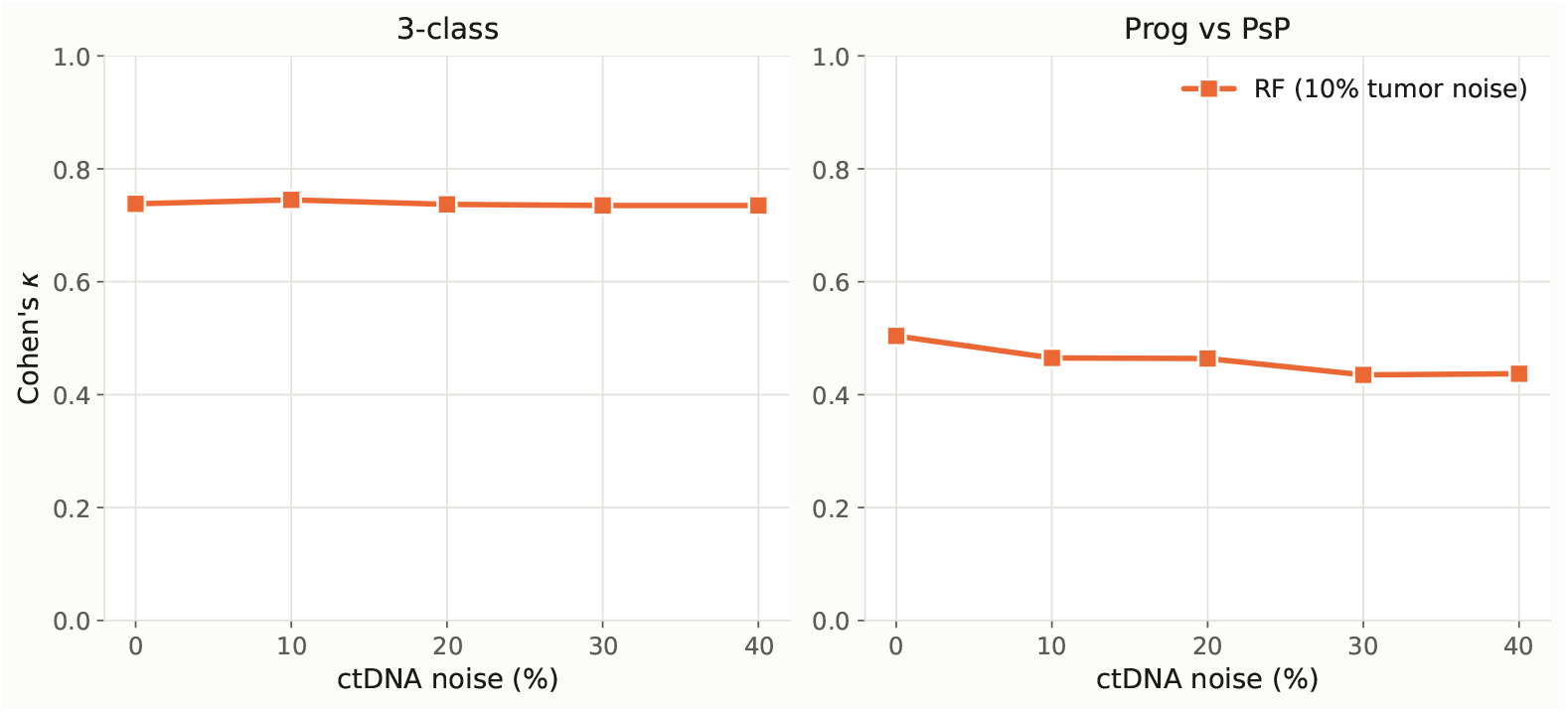
Cohen’s Kappa for classification of *n* = 9,934 *in silico* trajectories the random forest classifier (RF) with varied levels of lognormal ctDNA measurement noise added. Performance is analyzed for the 3 category classification with progression, pseudoprogression, and response/stable disease as well as for distinguishing progression and pseudoprogression. Constant lognormal tumor diameter measurement noise is added with 10% coefficient of variation.

We also examined the effect of reducing the ctDNA sampling window. The tumor sampling was unchanged with intervals of 25-50 days. We compared the original ctDNA sampling window of 25-50 days, with a halved window, a quartered window, and full daily sampling (Fig. 8). The tumor measurement noise was kept at 10% and the ctDNA measurement was 30% for all runs. Shortening the sampling window did not have a pronounced effect on the RF performance.

**Figure 8.**
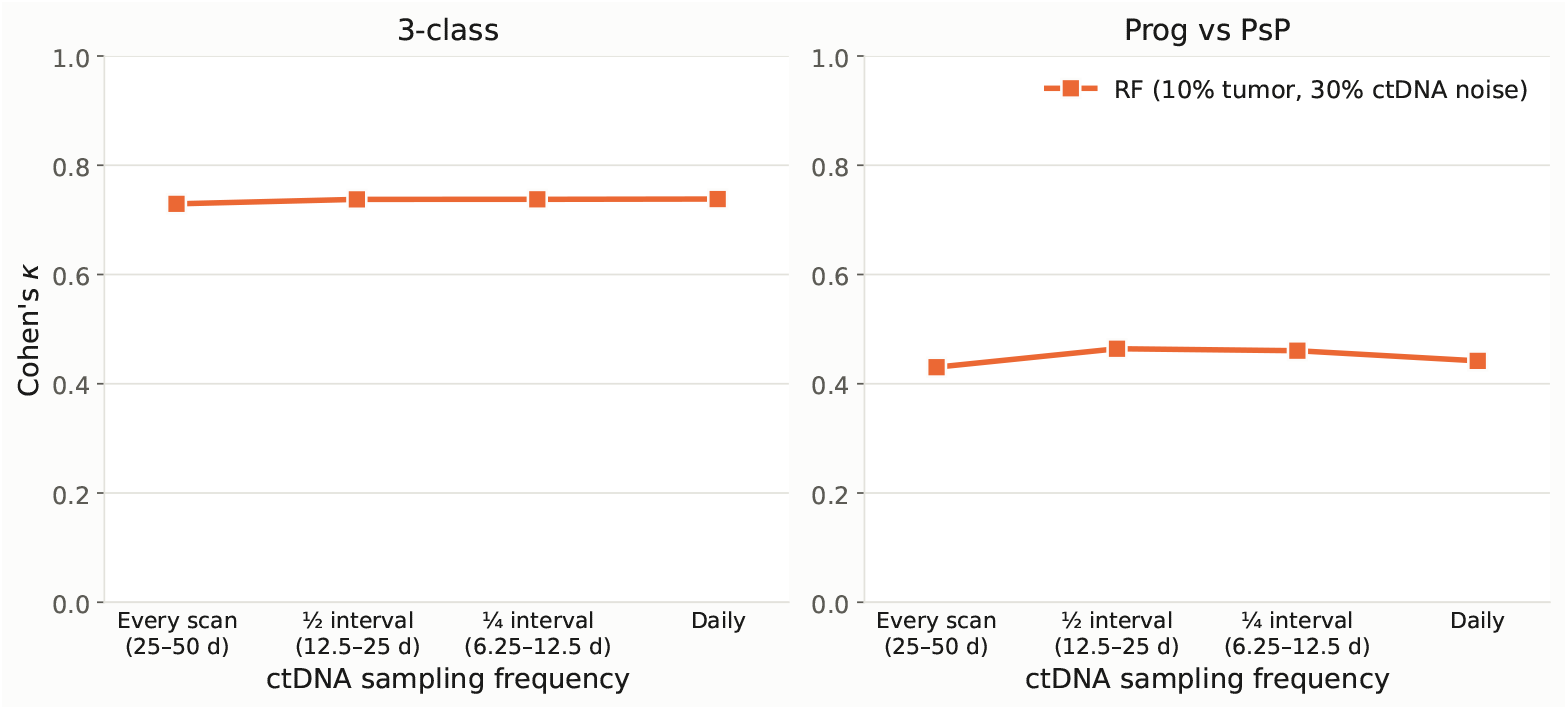
Cohen’s Kappa for classification of *n* = 9,934 *in silico* trajectories the random forest classifier (RF) with varied ctDNA sampling windows. The tumor measurement window is 25-50 days for all runs, while the ctDNA measurement window is varied. Performance is analyzed for the 3 category classification with progression, pseudoprogression, and response/stable disease as well as for distinguishing progression and pseudoprogression. Constant lognormal tumor diameter and ctDNA measurement noise is added with 10% and 30% coefficient of variation, respectively.

#### 2.4.3 Comparison with RECIST and iRECIST

Finally, we benchmarked our RF classifier by comparing it to clinical standards RECIST 1.1 and iRECIST. Recall that both RECIST 1.1 and iRECIST are primarily based on tumor measurements and do not incorporate ctDNA for classification. When an increase of 70% in tumor volume is observed, RECIST 1.1 immediately classifies the response as progression while iRECIST requires an additional follow-up observation before making its classification. Our RF classifier has access to both tumor and ctDNA data, but is restricted to the shorter window used by RECIST 1.1.

We compare RF, RECIST 1.1, and iRECIST across varied levels of tumor diameter measurement noise (Fig. 9). 30% ctDNA measurement noise is also added throughout. In our simulated cohort, for both tasks our RF outperforms RECIST 1.1 across all levels of tumor measurement noise. In the no-noise scenario, iRECIST performs better than the RF at distinguishing progression and pseudoprogression and marginally better at the 3-class task. However, under the modeled conditions the performance of the RF appears more robust to tumor measurement noise and improves upon iRECIST at moderate levels of measurement noise.

**Figure 9.**
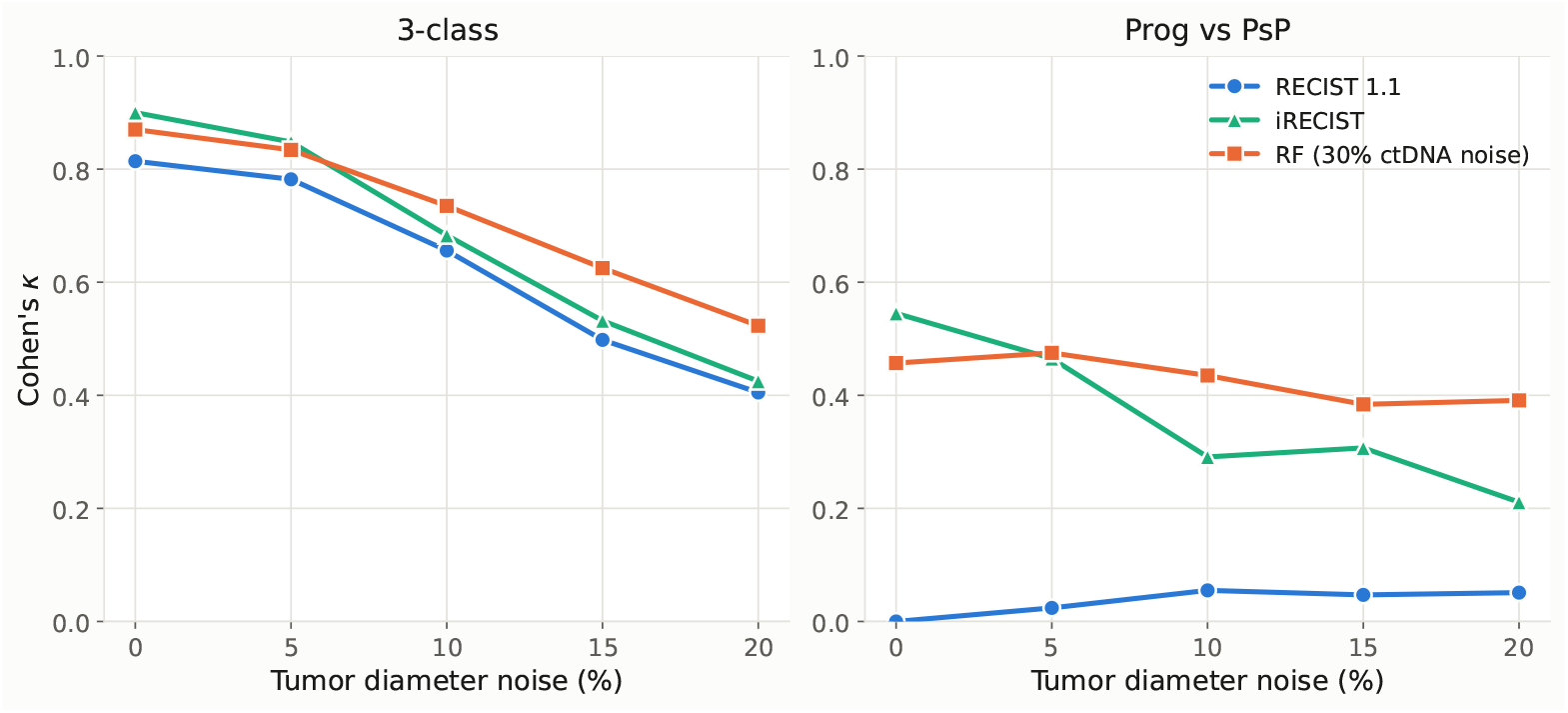
Cohen’s Kappa for classification of *n* = 9,934 *in silico* trajectories by RECIST 1.1, our random forest classifier (RF), and iRECIST with varied levels of lognormal tumor diameter measurement noise added. Performance is analyzed for the 3 category classification with progression, pseudoprogression, and response/stable disease as well as for distinguishing progression and pseudo-progression. For the RF classifier, constant lognormal ctDNA noise is added with 30% coefficient of variation.

Since iRECIST relies on a follow-up measurement to confirm the response class, our RF classifier was able to provide a classification a median of 37.5 days earlier than iRECIST. The faster classification provided by RF results in a substantial benefit of limiting additional growth in true progressors. Fig. 10 shows a histogram of the percent volume increases of true progressors between the time that progression is first observed and the follow-up assessment by iRECIST. The median additional growth was +144% in that time.

**Figure 10.**
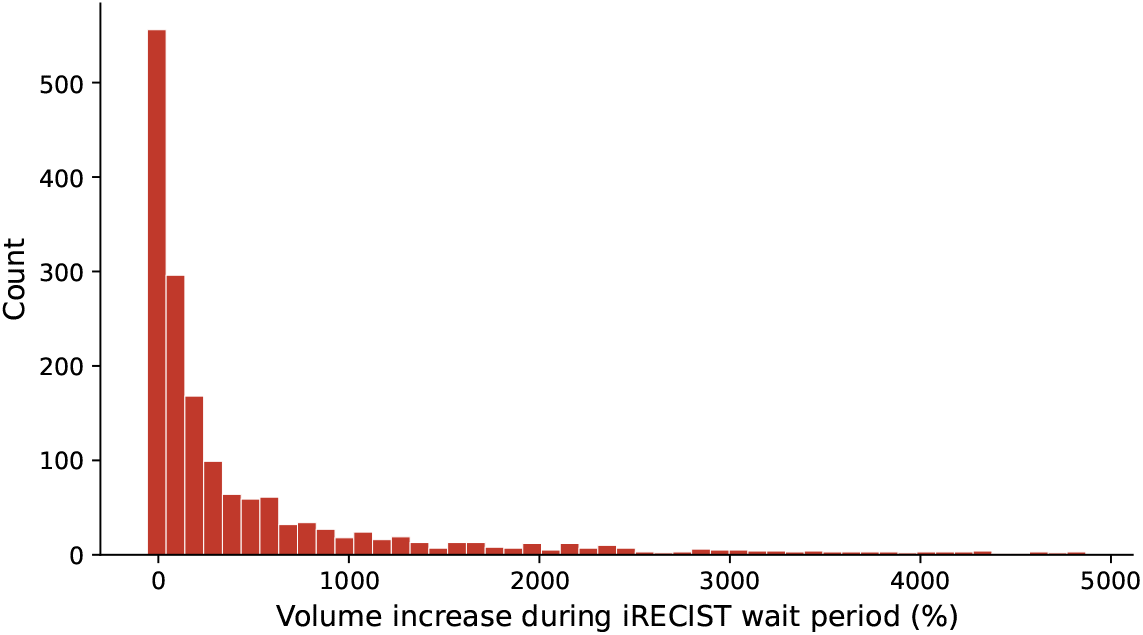
The *in silico* cohort with sampling noise included 1,755 patients categorized as true progressors. After initial observation of progression, iRECIST requires a subsequent follow-up observation to confirm progression. This delay resulted in a median wait time of 37.5 days and a median additional volume growth of +144% in true progressors.

The results of the comparison between RF, RECIST 1.1, and iRECIST are summarized in Table 3 for the cohort with 10% tumor measurement noise and 30% ctDNA measurement noise.

**Table 3.** Summary of RF, RECIST 1.1, and iRECIST classification performance in cohort with 10% tumor measurement noise and 30% ctDNA measurement noise.

|  | RF classifier | RECIST 1.1 | iRECIST |
| --- | --- | --- | --- |
| Data type | Tumor and ctDNA | Tumor | Tumor |
| Median days to classify progression | 45.7 | 44.3 | 82.1 |
| 3-class $\kappa$ | 0.735 | 0.656 | 0.683 |
| Prog. vs. PsP $\kappa$ | 0.435 | 0.055 | 0.291 |

#### 2.4.4 Validation of RF classifier on clinical data

Our ability to validate the performance of our random forest classifier outside of model-generated cohorts was limited by a lack of publicly available ICI treatment datasets with longitudinal tumor and ctDNA measurements. Bratman et al. [12] collected longitudinal tumor and ctDNA samples in solid tumor patients treated with pembrolizumab. Of the *n* = 23 patients with sufficient longitudinal tumor and ctDNA data samples, 3 trajectories were classified as progression and 20 were classified as response/stable disease according to our ground truth classification. The data set did not include any patients that would meet the criteria for pseudoprogression: an observed increase of at least 20% in the sum of long diameters (SLD), followed by control or decrease in SLD. Thus, this analysis evaluates classification of response/ stable disease vs progression, but does not provide clinical validation of pseudoprogression classification. Our RF classifier was able to correctly classify 3/3 progression trajectories and 19/20 response/stable disease trajectories. The single response/stable disease patient misclassified as progression by the RF classifier exhibited a tumor volume increase of 58.9% (16.7% increase in SLD). This is just under the threshold necessary for the ground truth to be classified as progression. Since this dataset did not include pseudoprogression, both RECIST and iRECIST were completely accurate by definition. In the absence of pseudoprogression, the RECIST and iRECIST criteria line up with our ground truth classification exactly. The absence of PsP cases in this cohort highlights the broader limitation of available datasets to accurately determine whether or not ctDNA can help resolve ambiguities in assessing treatment response due to PsP. To do this would require paired measurements collected specifically across the period of suspected progression and confirmation of response at follow-up. Largeer clinical datasets, containing pseudoprogression cases, will be required to determine whether the performance improvements observed in our model-generated *in silico* cohort generalize to clinical settings.

## 3 Discussion

We introduce a novel mathematical model of tumor-immune dynamics under ICI treatment. The terms governing the tumor-immune interactions and state transitions are well supported by fundamental mathematical models [23–25] and scientific literature [2, 26–29]. In our model, we define the measured tumor volume *V* = *T* + *I*^*A*^ + *I*^*E*^ to include both tumor and immune cells, and explore ctDNA shedding in the context of ICI treatment. Our model closely reproduces clinical patient trajectories (Section 2.2) and achieves higher *R*^2^ and lower MAE than models introduced in [15], albeit with more parameters.

Using our mechanistic model, we demonstrate how pseudoprogression dynamics can naturally arise from simple biological principles. We present an analytic perspective on the definition of pseudoprogression (Fig. 3) and compare the analytic and clinical classifications of patient response (Fig. 4). In our *in silico* cohort, analytical pseudoprogressors did exhibit transient growth followed by disease response, but the majority did not meet the threshold for clinical classification of pseudoprogression: an increase of 20% in SLD (≈70% in volume). Our simulations show that transient growth trajectories, consistent with the underlying mechanisms of pseudoprogression, may occur more frequently than trajectories meeting the clinically-defined PsP criteria. Thus the incidence of PsP [4] may depend partly on clinical sampling times and measurement resolution, and warrants further evaluation in clinical datasets. Our model also provides insight into potential mechanisms of pseudoprogression; in particular, initial tumor growth due to delayed immune activity and immune infiltration may both contribute to the transient increase in measured tumor size.

The classification of pseudoprogression presents a complex clinical challenge: when tumor growth is observed, should it be classified as progression immediately or should treatment continue until the next assessment? Under RECIST 1.1 [7] pseudoprogression may be miscategorized as progression, resulting in premature cessation of potentially effective treatment. Under iRECIST [6], waiting for follow-up assessment may benefit pseudoprogressors, but can cause unnecessary delays in pivoting treatment plans in true progressors. We therefore asked whether ctDNA dynamics could be integrated with tumor dynamics to improve classification without relying on follow-up assessments. Using Cohen’s Kappa [38] to assess classification of progression, pseudoprogression, and response/stable disease in a model-generated *in silico* cohort, we found that a random forest classifier using ctDNA and tumor dynamics performed better than iRECIST in the presence of tumor measurement noise, and performed significantly better than RECIST 1.1 in the scenarios considered (Fig. 9). We hypothesize that the rigid thresholds of iRECIST and RECIST 1.1 make them more susceptible to the influence of noise. Moreover, by delivering the prediction a median of 37.5 days earlier than iRECIST, the RF classifier was able to avoid a median of +144% additional growth from true progressors (Fig. 10). Note also that 37.5 days is near the lower end of the 4-8 week follow-up window recommended by iRECIST, with the median as high as 6.8 weeks in some studies [39]. These simulations help quantify the tradeoffs of delayed classification, as it can protect patients from premature treatment discontinuation while also prolong ineffective therapy for patients with true progression.

Our work provides insight into the dynamics and mechanisms underlying pseudoprogression in ICI treatment and demonstrates the predictive potential of model-informed analysis of paired tumor and ctDNA data. In our simulated cohorts, when comparing RF classifiers using tumor data only, ctDNA data only, and coupled tumor-ctDNA data, the RF with access to coupled data performed best in all scenarios (Fig. 6). Properly harnessing the combined signal may improve timely and accurate categorization of patient response. Variability in ctDNA shedding and measurement can result in an unfavorable signal-to-noise ratio [40]. However, our RF classifier is largely able to maintain consistent accuracy across a range of ctDNA measurement noise levels (Fig. 7). Moreover, our RF does not rely on additional ctDNA sampling times beyond the tumor evaluation schedule (Fig. 8), which would reduce the impact of ctDNA sampling on patients. Our previous work on early ctDNA sampling [21], focused on initial evaluation of potential treatment benefit and thus relied on high frequency ctDNA sampling within the first days of treatment to capture transient peaks. The timescale of identifying and predicting pseudoprogression is on the order of weeks or months, rather than days, so the transient dynamics are not essential to our RF performance.

While we have grounded our analysis with scientific literature and validation on publicly available data, this work is limited by the lack of adequate longitudinal tumor and ctDNA data sets in patients receiving ICI treatment, combined with the relatively low incidence of clinically-characterized pseudoprogression. Definitive evaluation of early PsP discrimination will require prospective (or carefully curated retrospective) cohorts with synchronous serial imaging and ctDNA measurements covering the time period of initial treatment through confirmation of progression or treatment response.

Possible directions for further exploration include modeling the evolution of treatment resistance, quantifying the impact of measurement noise, examining peripheral immune activity data, and modeling of Chimeric antigen receptor (CAR)-T-cell therapy. Our model currently does not account for the evolution of treatment resistance within the tumor. Our group has previously used a stochastic branching processes to model the evolution of resistance to targeted therapy via point mutation or gene amplification [41] and a similar framework could be applied to tumor-immune dynamics under ICI treatment. Peripheral blood lymphocyte [42] and Ki67+CD8+ T cell [43] counts before and during treatment may help predict response to ICI treatment. We briefly explored the effect of including the baseline immune level *y*_0_ and longitudinal *y* measurements on the tumor-ctDNA random forest classifier’s performance in Fig. A1. The inclusion of *y*_0_ offered no improvement over the RF classifier, but including longitudinal *y* measurements did provide a modest improvement. Pseudoprogression has also been observed during CAR-T-cell therapy [44]. In Appendix C, we briefly demonstrate how our model provides a flexible framework that can naturally be extended to model the infusion of CAR-T cells, but further exploration is needed. The dynamics of our ICI model generally result in the immune population being capped by the asymptote *y*_∞_ (Appendix A). However, CAR-T infusion could allow the immune population to exceed that asymptote and necessitate the analysis of the previously infeasible upper regions of the phase plane.

In summary, our results provide a mechanistic framework linking ctDNA dynamics with tumor growth, immune infiltration and treatment response during ICI therapy. The model predicts that paired ctDNA and tumor burden longitudinal dynamics may enable earlier determination of pseudoprogression vs. true progression. Prospective collection of tumor imaging and ctDNA measurements until confirmation of response or true progression is now needed to test whether this modeling prediction can be translated into improved response assessment in the clinic.

## 4 Methods

### 4.1 Analytic exploration

Examining just the tumor cells and the active immune cells and fixing *f* (*t*) = 1, we have

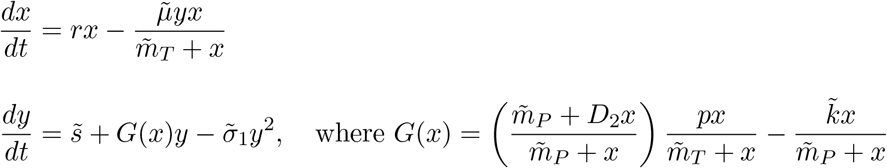

The tumor (*n*_*x*_), immune (*n*_*y*_), and total (*n*_*x*+*y*_)nullclines of the system are equations that are satisfied by points where 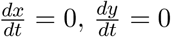, and 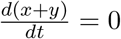, respectively. Following derivations shown in Appendix A, we have:

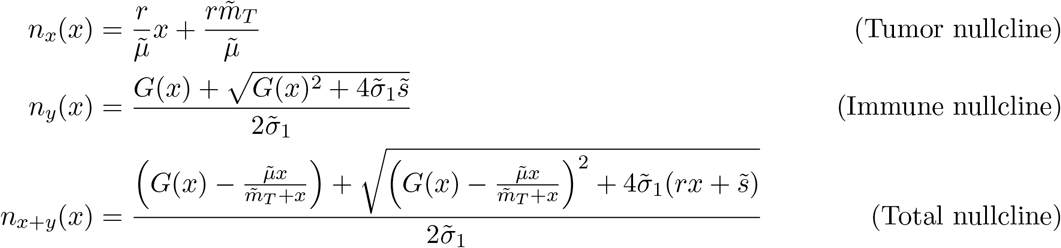

Deriving a closed form of the separatrix *S* is not tractable, but we can derive its form numerically using `scipy.optimize.fsolve` to locate the saddle point and `scipy.integrate.solve_ivp` (RK45) to backward-integrate the stable manifold.

### 4.2 Model fitting

Each patient’s longitudinal data were normalized to baseline before fitting, so that *V* (0) = 1 and *C*(0) = 1. Changes in SLD were converted to changes in volume by assuming spherical tumors. Since our focus was on initial response patterns, we excluded delayed progression that arose after initial patient response.

The four-state ODE system (tumor *x*, active immune *y*, exhausted immune *z*, and ctDNA *C*) was simultaneously fitted to the observed *V* (*t*) = *x* + *y* + *z* and *C*(*t*) trajectories for each patient. The objective was the sum of squared residuals

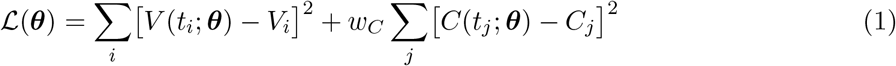

Analyzing the tumor and ctDNA data in Bratman et al. [12], we found that the variance of the ctDNA data was higher than variance of the tumor data by a factor of 3.1. Thus, we set the error weight of the ctDNA data to *w*_*C*_ = 0.3 in order to account for the larger measurement variability of the ctDNA signal.

Ten free parameters were estimated per patient (Table 4). Three parameters with low variance in initial fitting were fixed at population-representative values 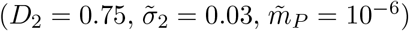. The ctDNA clearance rate was fixed at *ε* = 25 d^−1^, corresponding to a plasma half-life of approximately 40 minutes, consistent with published ctDNA kinetics [37]. The ctDNA shedding coefficient *α* was derived analytically from the normalization constraint *C*(0) = 1 under the approximating assumption of no-drug steady-state conditions similar to those used in [16]:

**Table 4.** Free parameters estimated per patient, with optimization search bounds. Parameters marked *†* are searched on a log_10_ scale.

| Parameter | Description | Lower | Upper |
| --- | --- | --- | --- |
| $r$ | Tumor net proliferation rate [45] | $10^{-3}$ | 0.10 |
| $\tilde{\mu}$ | Net immune killing rate | 0.005 | 1.00 |
| $\tilde{m}_T$ | Scaled tumor saturation constant$^\dagger$ | $10^{-6}$ | 1.00 |
| $\tilde{s}$ | Scaled immune recruitment rate$^\dagger$ | $10^{-8}$ | $10^{-2}$ |
| $p$ | Immune proliferation rate | 0.01 | 1.00 |
| $\tilde{\sigma}_1$ | Scaled immune competition rate$^\dagger$ | $10^{-3}$ | 50.0 |
| $\tilde{k}$ | Net immune exhaustion rate | $2.5 \times 10^{-4}$ | 0.25 |
| $x_0$ | Initial normalized tumor volume | 0.50 | 0.995 |
| $y_{\text{frac}}$ | Initial active immune fraction | $10^{-3}$ | 0.999 |
| $\phi$ | Intrinsic death fraction | 0.05 | 0.99 |

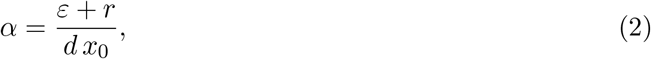

where *d* = *rϕ/*(1 − *ϕ*) is the intrinsic tumor death rate. The initial immune compartment was parameterized through *y*_frac_, the fraction of total non-tumor volume occupied by active immune cells at *t* = 0, with *y*_0_ = (1 − *x*_0_) *y*_frac_ clamped to 2 *y*_∞_ to prevent physiologically implausible initial conditions.

Optimization used bounded nonlinear least squares (`scipy.optimize.least squares`, trust-region reflective method) with 10 random restarts drawn uniformly from the search bounds (Table 4), and an additional 70 restarts for patients whose best fit achieved *R*^2^ *<* 0.80.

#### Tumor-only data sets

For the tumor-only POPLAR [33], BIRCH [34], OAK [35], and FIR [36] data sets, the ctDNA equation was omitted and the fitting reduced to a three-state (*x, y, z*) ODE against normalized tumor volume alone. Accordingly, the parameters *ϕ, ε, w*_*C*_, and *α* were not applicable, reducing the free parameter count to nine.

#### *in silico* cohort

We generated a cohort of 10,000 virtual patients by forward-simulating the three-state ODE (tumor *x*, active immune *y*, exhausted immune *z*) over a 300-day horizon with daily time steps. Empirical parameter distributions were derived by fitting the model to the POPLAR [33], BIRCH [34], OAK [35], and FIR [36] data sets (Table 1). Parameters for each virtual patient were drawn jointly from a Gaussian copula whose marginals were the empirical distributions of the fitted parameter values and whose dependence structure was the rank correlation matrix of those fits. In Appendix A, we derive an immune saturation limit *y*_∞_. In order to avoid unrealistic initial immune fractions, draws violating *y*_0_ ≤ 2*y*_∞_ were rejected and redrawn. The intrinsic death fraction *ϕ* was sampled uniformly on [0.20, 0.90] and used only for the ctDNA equation (*ε* = 25 d^−1^ fixed), which was simulated alongside the ODE to produce synthetic ctDNA trajectories. All three fixed model parameters 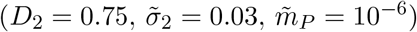 were held at their population values.

To generate *in silico* clinical data sets from these simulations, the daily trajectories were sampled according to a clinical observation schedule. For each patient, tumor (*V*) and ctDNA (*C*) values were sampled at baseline (*t* = 0) and for subsequent samples, the inter-visit intervals were drawn uniformly from 25–50 days. Cohorts were generated with varied levels of lognormal measurement noise applied to the *V* and *C* values. The tumor diameter noise ranged from 0 to 20% coefficient of variation and the ctDNA noise ranged from 0 to 40%.

Ground truth clinical response labels were applied to each patient based on the trajectory exhibited by the data on the clinical sample schedule. Since a follow-up observation is required to distinguish pseudoprogression and response/stable disease, we excluded 66 patients who exhibited progression near the 300-day time horizon and did not receive a follow-up measurement. This resulted in a cohort size of *n* = 9,934 patients in total.

### 4.3 Random forest classifiers

We examined three random forest classifiers: RF (all), RF (ctDNA), and RF (tumor). RF (all) had access to *V* and *C* data, RF (ctDNA) was restricted to *C* only, and RF (tumor) was restricted to *V* only. Each classifier received time series data up until the first sample time where progression was observed. That is, the classifiers operate on the same sampling schedule as RECIST 1.1 and have access to one fewer sampling time than iRECIST. Each patient’s observation schedule was drawn independently, with inter-visit intervals sampled uniformly from 25–50 days starting from *t* = 0, yielding 8–9 observations per patient over the 300 day window (mean interval ≈ 37 d). The feature vector comprised the observation times together with the corresponding tumor volumes, ctDNA values, or both, padded to a fixed length of 14 slots. Slots beyond the observed progression time or the end of the window were set to `NaN`. The forest comprised 300 trees with balanced class weights to account for the class imbalance between progression and pseudoprogression patients. All other hyperparameters were left at their `scikit-learn` defaults. For each level of noise, the corresponding dataset was split into 70% training and 30% test sets using a stratified random split (seed 42), and all rule-based comparators (RECIST 1.1, iRECIST) were evaluated on the same held-out test set.

## 5 Code availability

All model code, including ODE implementation, parameter fitting, forward simulations, and RECIST comparison scripts, is available on Zenodo at https://doi.org/10.5281/zenodo.21345147.

## Acknowledgments

J.F. was partially supported by NSF DMS 2052465 and NSF CMMI 2228034, the Rainwater Foundation and the Ellison Institute. A.L. was partially supported by NSF DMS 2052465. K.L. was partially supported by NSF CMMI 2228034. J.F would like to thank Bo Connelly for useful initial discussions on this topic.

## A Example heuristic

Notice that 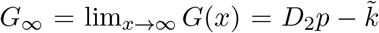. Then the immune saturation limit (created by the immune competition) is given by the asymptote

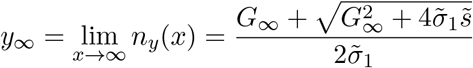

Notice that the tumor nullcline *n*_*x*_(*x*) intersects this asymptote at 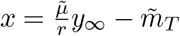. Now consider (*x*_0_, *y*_0_) such that 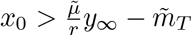 and *y*_0_ ≤ *y*_∞_. Then

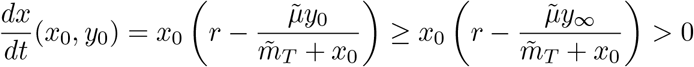

So points in this region will tend toward tumor escape. Moreover, notice that as 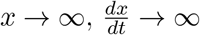 while 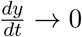 so heuristically 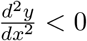 for *x* sufficiently large and *y* sufficiently close to *y*_∞_. In other words, trajectories here will eventually be concave down and “curve to the right.” Detection of this signal in longitudinal data may be a path toward identification of true progression.

## B Progression versus pseudoprogression classification with immune measurements

**Figure A1.**
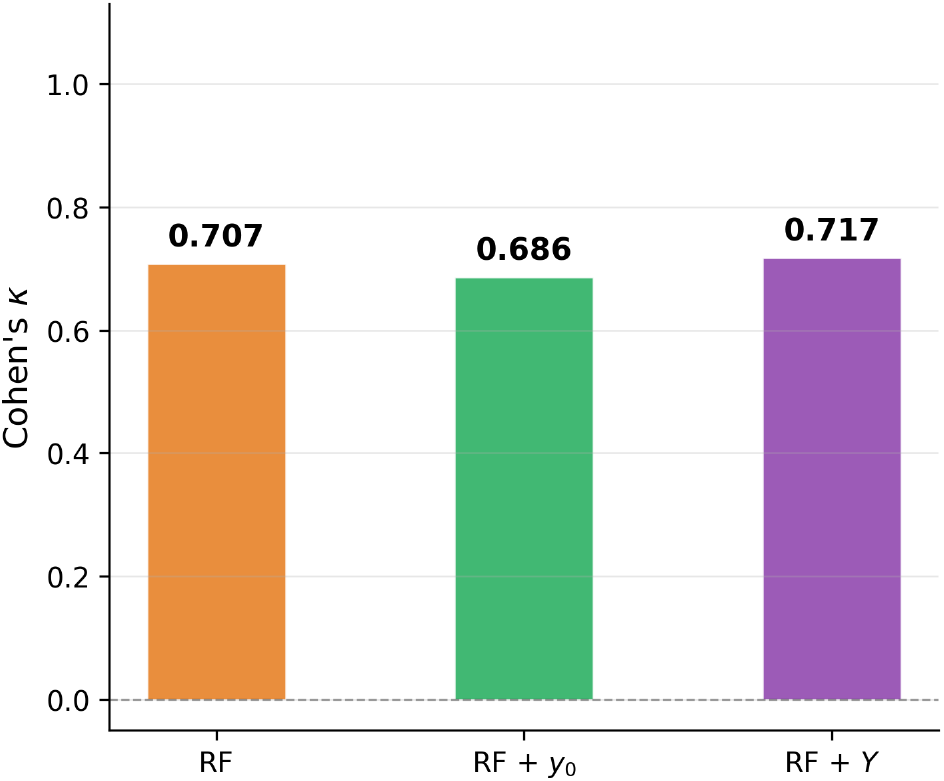
Progression vs. pseudoprogression classification with immune measurements. We examine the effect of incorporating an initial baseline immune measurement (*y*_0_) and longitudinal immune measurements (*Y*) on the performance of our random forest classifier (RF). The baseline immune measurement did not improve the Cohen’s *κ* of the classifier when attempting to distinguish progression and pseudoprogression. Longitudinal immune measurements provide a modest improvement.

## C CAR-T modeling

Chimeric antigen receptor (CAR)-T-cell therapy involves the infusion of genetically engineered T cells with improved anti-tumor behavior [46]. Our model can easily be extended to model CAR-T therapy as follows:

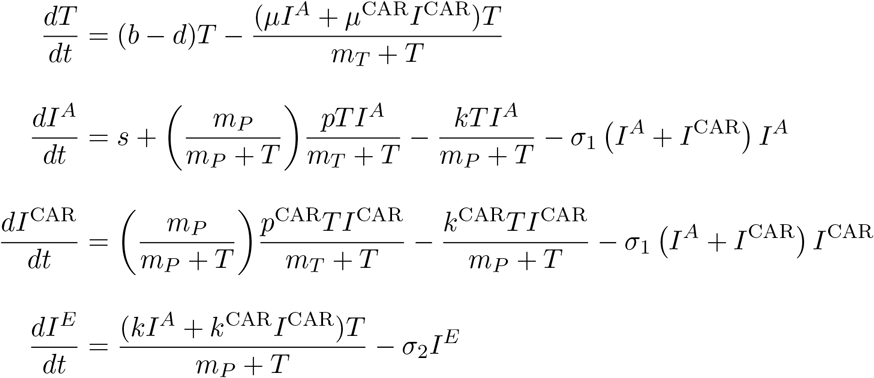

We include an additional compartment *I*^CAR^ to represent the CAR-T cells and omit the ICI concentration terms involving *f* (*t*). The CAR-T cells have their own tumor-killing, proliferation, and exhaustion coefficients *µ*^CAR^, *p*^CAR^, *k*^CAR^. Note that the lymphocyte recruitment term *s* is not included for the CAR-T cells because the only source of CAR-T cells is the initial infusion and CAR-T proliferation. We model the infusion by setting *I*^CAR^(0) high. In our exploration, the immune population is generally limited by the immune saturation asymptote *y*_∞_ (Appendix A). However, the CAR-T infusion may allow the initial immune population to exceed that bound, opening up exploration of the upper regions of the phase plane that were not feasible to reach in the ICI regime.

## Notes

### Competing Interest Statement

The authors have declared no competing interest.

